# Cannabidiol attenuates seizures and EEG abnormalities in Angelman syndrome model mice

**DOI:** 10.1101/689943

**Authors:** Bin Gu, Manhua Zhu, Madison R. Glass, Marie Rougié, Viktoriya D. Nikolova, Sheryl S. Moy, Paul R. Carney, Benjamin D. Philpot

**Author notes:** Address correspondence to: Ben Philpot, University of North Carolina, 116 Manning Dr, Chapel Hill, NC 27599.

## Abstract

Angelman syndrome (AS) is a neurodevelopmental disorder characterized by intellectual disability, lack of speech, ataxia, EEG abnormalities, and epilepsy. Seizures in AS individuals are often refractory to existing antiepileptic medications. Therefore, there is an unmet need for better seizure control, which could potentially improve other symptomatic domains such as cognitive function. Cannabidiol (CBD), a major phytocannabinoid constituent of cannabis, has anti-seizure activity and behavioral benefits in preclinical and clinical studies for some disorders associated with epilepsy, suggesting that the same could be true for AS. Here we show that acute CBD (100 mg/kg) attenuated hyperthermia- and acoustically-induced seizures in a mouse model of AS. However, neither acute CBD nor a two-weeklong course of CBD administered immediately after a kindling protocol could halt the pro-epileptogenic plasticity observed in AS model mice. CBD had a mild sedative effect, but did not have a major impact on motor performance. CBD abrogated the enhanced delta rhythms observed in AS model mice, indicating that CBD administration could also help normalize the EEG deficits observed in individuals with AS. Our results provide critical preclinical evidence supporting CBD treatment of seizures and alleviation of EEG abnormalities in AS, and will thus help guide the rational development of CBD as an AS adjunctive treatment.

## INTRODUCTION

Deletions or mutations of the maternally inherited copy of the *UBE3A* gene cause Angelman syndrome (AS). Individuals with AS exhibit developmental delay, motor dysfunction, minimal speech, highly penetrant EEG abnormalities, and seizures (1, 2). Epilepsy in AS is common (80%–95%), polymorphic, and often resistant to available antiepileptics. The frequency, severity, and pharmacological intractability of the seizures exacts a heavy toll on individuals with AS and their caregivers (3-6). AS model mice lacking a functional maternal copy of the orthologous *Ube3a* gene (*Ube3a*^*m–/p+*^) phenocopy many clinical aspects of AS, including seizure susceptibility and EEG abnormalities, thereby offering a preclinical model for developing new therapeutics (7-15).

Cannabidiol (CBD), a major phytocannabinoid constituent of cannabis, is gaining attention for its antiepileptic, anxiolytic, and antipsychotic properties (16). In 2018, the FDA approved CBD (Epidiolex^®^) for the treatment of seizures associated with two rare and severe forms of epilepsy — Lennox-Gastaut syndrome and Dravet syndrome. While the interest and off-label medical use of CBD has largely outpaced the preclinical and clinical research, CBD may provide a viable treatment for several other neurological disorders associated with epilepsy such as AS (17-20). However, little is known about the potential antiepileptic and other benefits of CBD in AS. The broad beneficial spectrum of CBD makes it a unique candidate for treating the complex symptoms associated with AS, as it holds the potential to simultaneously ameliorate behavioral deficits, EEG abnormalities, and seizures (21).

In this study, we systematically tested the beneficial effects of CBD on core AS-related pathologies in mice, with the expectation that this information will guide eventual clinical use. We report that acute treatment of CBD substantially attenuated audiogenic and hyperthermia-induced seizure severity, and normalized delta rhythms in AS model mice. The anticonvulsant dose of CBD (100 mg/kg) caused mild sedation, but had little effect on motor coordination or balance. While acute CBD could suppress seizure severity, CBD stopped short of being able to prevent the pro-epileptogenic plasticity observed in AS model mice. Our study provides a preclinical framework to better guide the rational development of CBD as an AS treatment.

## RESULTS & DISCUSSION

As with individuals with AS, mice with maternal loss of *Ube3a* exhibit epileptic phenotypes. For example, AS mice on a 129 background have elevated seizure responses to acoustic stimuli (9, 10, 14). We verified that AS model mice (129 background) exhibited more frequent audiogenic seizures than wild-type (WT) littermates and had a far greater likelihood of progressing from wild running to a more severe tonic-clonic episode (Figure 1). Acute treatment of CBD at higher doses significantly reduced the frequency (@ 100 mg/kg CBD) and severity (@ 50 and 100 mg/kg CBD) of audiogenic-induced seizures (Figure 1). These results suggest strong dose-dependent anticonvulsant effects of CBD in AS model mice.

**Figure 1.**
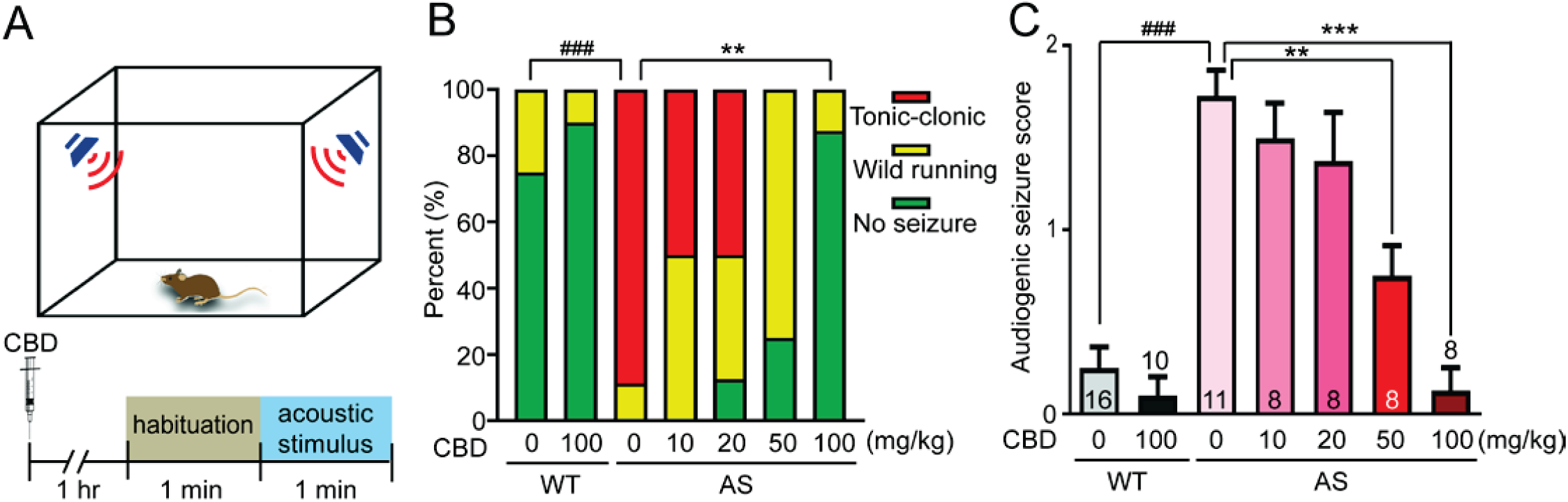
CBD attenuates audiogenic seizures in AS model mice. (A) Schematic of audiogenic-induced seizure paradigm. (B) Treatment (i.p.) of CBD at 100 mg/kg significantly reduced audiogenic-induced seizure frequency. n=8-16 mice/group. ### p<0.001 compared to WT-Veh; ** p<0.01 compared to AS-Veh, Fisher’s exact test. (C) CBD at 50 and 100 mg/kg significantly reduced audiogenic-induced seizure severity. Score 0 = no seizure response; Score 1 = wild running and jumping; Score 2 = tonic-clonic clonus. n=8-16 mice/group. ### p<0.001 compared to WT-Veh; ** p<0.01, ***p<0.001 compared to AS-Veh, two-way ANOVA with Tukey’s *post hoc* test.

We recently implemented the flurothyl kindling and retest paradigm in AS model mice (C57BL/6J background), and found that AS model mice responded to both initial (day1) seizure induction and kindling (day1 – day8) similarly to WT mice, but they displayed a markedly increased sensitivity to flurothyl-induced seizure measured a month later at retest (day36) (8). Elevation of core body temperature also triggered convulsions in kindled AS but not WT mice, resembling the clinical observation that AS individuals are susceptible to febrile seizures with moderate increases in body temperature (3, 4, 8). To test whether the anticonvulsant effects of CBD are generalizable across different seizure induction paradigms, we treated flurothyl-kindled mice acutely with CBD (100 mg/kg) 1 hr prior to the flurothyl- or hyperthermia-stimuli (Figure 2A). Surprisingly, the acute administration of CBD had little effect on flurothyl-induced seizure threshold (Figures 2B and 2C) or the body temperature for onset of hyperthermia-induced seizure in kindled AS mice (Figure 2D). However, CBD significantly attenuated the severity and duration of hyperthermia-induced seizures in kindled AS mice (Figures 2E and 2F), once again supporting a role for CBD in attenuating seizure severity.

**Figure 2.**
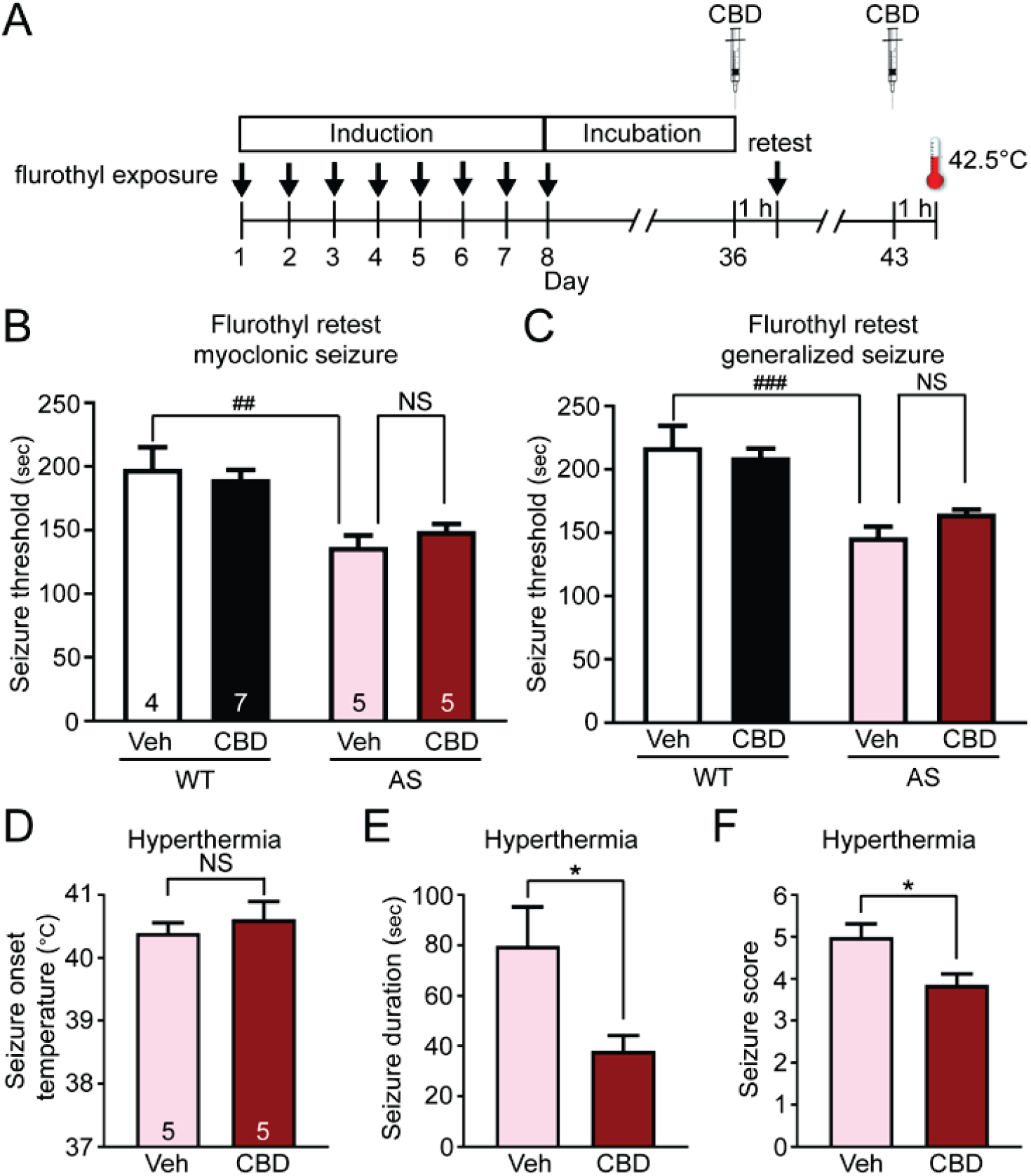
CBD reduces hyperthermia-induced seizure duration and severity in kindled AS model mice. (A) Schematic of flurothyl kindling and retest followed by hyperthermia-induced seizure. (B) Myoclonic and (C) generalized seizure threshold at flurothyl retest of kindled WT or AS model mice treated with either Veh or CBD (100 mg/kg, i.p.) 1hr prior to the retest. n=4-7 mice/group. ## p<0.01 and ### p<0.001 compared to WT-Veh, two-way ANOVA with Tukey’s *post hoc* test. Hyperthermia-induced seizure (D) onset body temperature, (E) duration, and (F) maximum seizure score of kindled AS mice treated with Veh or CBD (100 mg/kg, i.p.). n=5 mice/group. * p<0.01 compared to AS-Veh, unpaired t-test (two-tailed).

Aside from the profound anticonvulsant effect of CBD, little is known about the possible antiepileptogenic effect of CBD. This can be studied in a flurothyl kindling and retest model, as AS mice show similar seizure susceptibility to WT mice across an initial 8 days of flurothyl kindling, but then exhibit a pro-epileptogenic plasticity during the subsequent month-long incubation period that renders them highly susceptible to seizures compared to WT mice (8). To test whether CBD can exert an antiepileptogenic effect, we initiated a 2-week CBD treatment (100 mg/kg, i.p. once per day) immediately after completion of flurothyl kindling followed by a 2-week drug washout prior to the flurothyl retest (Supplemental Figure 1A). Consistent with our previous findings (8), vehicle-treated AS mice exhibited enhanced seizure susceptibility at flurothyl retest compared to WT mice (Supplemental Figures 1B and 1C). Moreover, the post-kindling CBD treatments failed to attenuate the enhanced seizure susceptibility measured in AS model mice at flurothyl retest (Supplemental Figures 1D and 1E), suggesting that two-week long CBD treatments (at 100 mg/kg) cannot prevent the pro-epileptogenic plasticity that occurs following kindling in AS mice.

Motor and behavioral impairments are common in AS children, significantly impact their daily lives, and increase the burden of their caregivers. No effective drug treatments are available. AS model mice exhibit behavioral and motor deficits (7, 9, 15, 22), some of which resemble those observed in AS individuals. To explore possible behavioral benefits of CBD, we treated AS model mice and WT controls with various doses of CBD 1 hr prior to behavioral testing. Similar to prior observations (7, 9, 15, 22), we found that vehicle-treated AS model mice exhibit impaired locomotor activity, poor motor coordination, and reduced marble burying behavior (Figure 3). Open field activity was reduced in AS model mice, and CBD had a dose-dependent sedative effect in both WT and AS mice (Figures 3A and 3B). Consistent with previous findings in rats (23), CBD did not have a major impact on motor skill learning and memory, regardless of genotype, as measured in rotarod acquisition and retest (Figures 3C and 3D). Notably, CBD at 100 mg/kg exaggerated the marble burying deficits of AS model mice (Figure 3E), which might be a consequence of its sedative effects.

**Figure 3.**
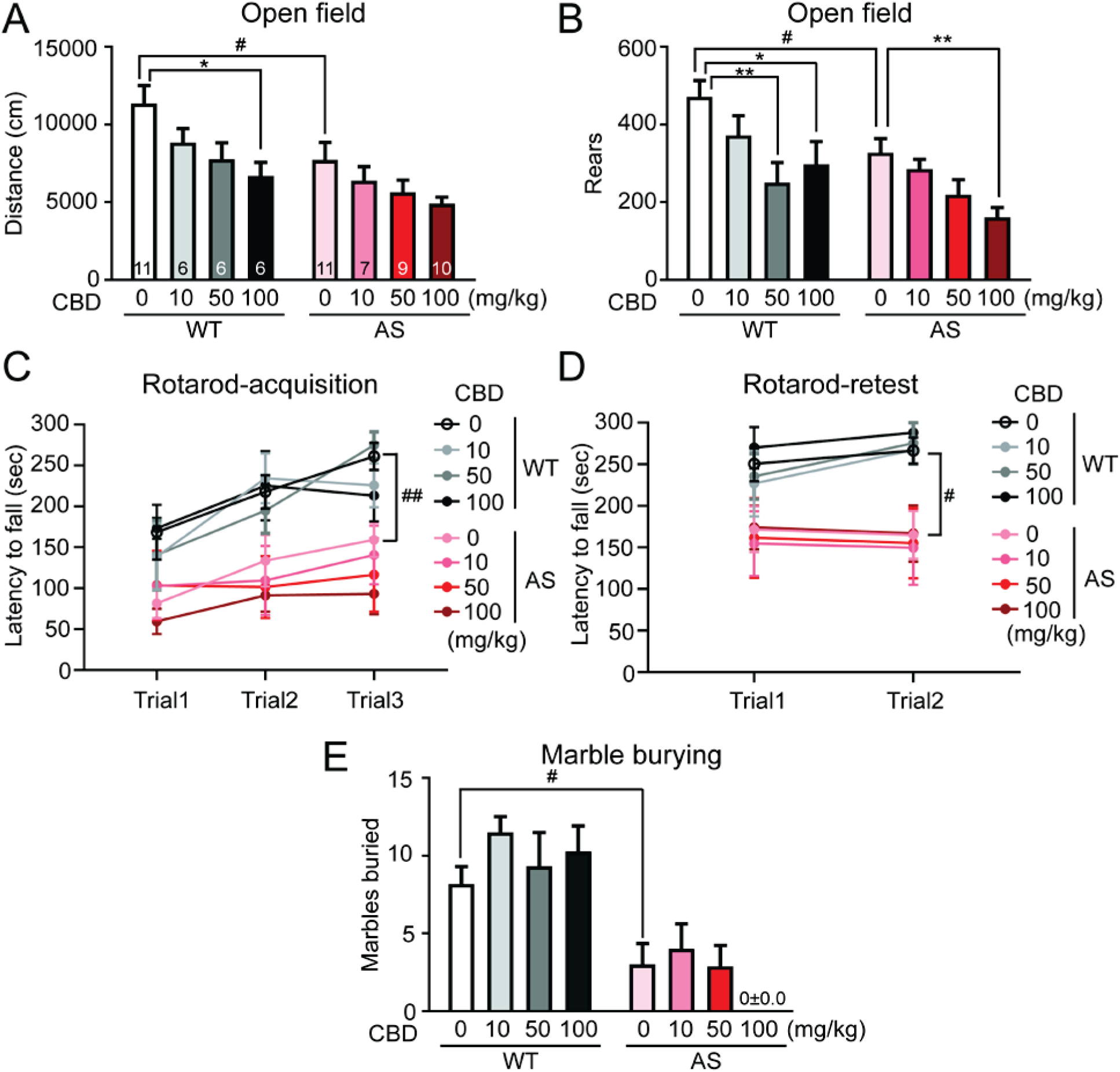
CBD exhibits moderate sedative effects in AS model mice. (A and B) WT and AS model mice were tested in open field to assess (A) horizontal (distance traveled) and (B) vertical (rears) movement 1 hour after Veh or CBD injection (i.p.). (C and D) Latency to fall from an accelerating rotarod in each trial of (C) acquisition and (D) retest session. Veh or CBD was injected (i.p.) 1 hour prior to acquisition session. (E) Number of marbles buried by WT and AS model mice treated with Veh or CBD (i.p.) 1 hour prior to the test. Note that none of the 10 AS model mice treated with 100 mg/kg CBD buried a marble during the test. n=6-10 mice/group. # p<0.05 and ## p<0.01 compared to WT-Veh; * p<0.05, **p<0.01 compared to AS-Veh, two-way ANOVA with Tukey’s *post hoc* test.

AS individuals have higher EEG power across all frequencies compared to neurotypical controls, with the largest difference manifested in a prominent peak in the delta frequency range (13, 24-26). Electrophysiological recordings from freely roaming AS model mice suggest strain- and region-dependent quantitative differences in EEG power spectrum analysis — the most robust elevations of delta and theta power were found in cortex of AS mice on a C57BL/6J background regardless of light cycle (7). Here we monitored and quantified cortical local field potentials (LFPs) of freely roaming AS and WT mice (C57BL/6J) after a 2-week vehicle or CBD treatment (Figure 4). Consistent with previous findings (7), AS model mice exhibited enhanced electrophysiological power compared to WT mice, particularly in delta (1–4 Hz) and theta (5–8 Hz) activity. CBD treatments had little effect on LFP power in WT mice, whereas the treatment significantly lowered total LFP power, including both delta and theta activity in AS model mice, normalizing levels to those observed in WT mice (Figure 4). All mice showed similar weight gain over the two weeks regardless of genotype and treatment (Supplemental Figure 2).

**Figure 4.**
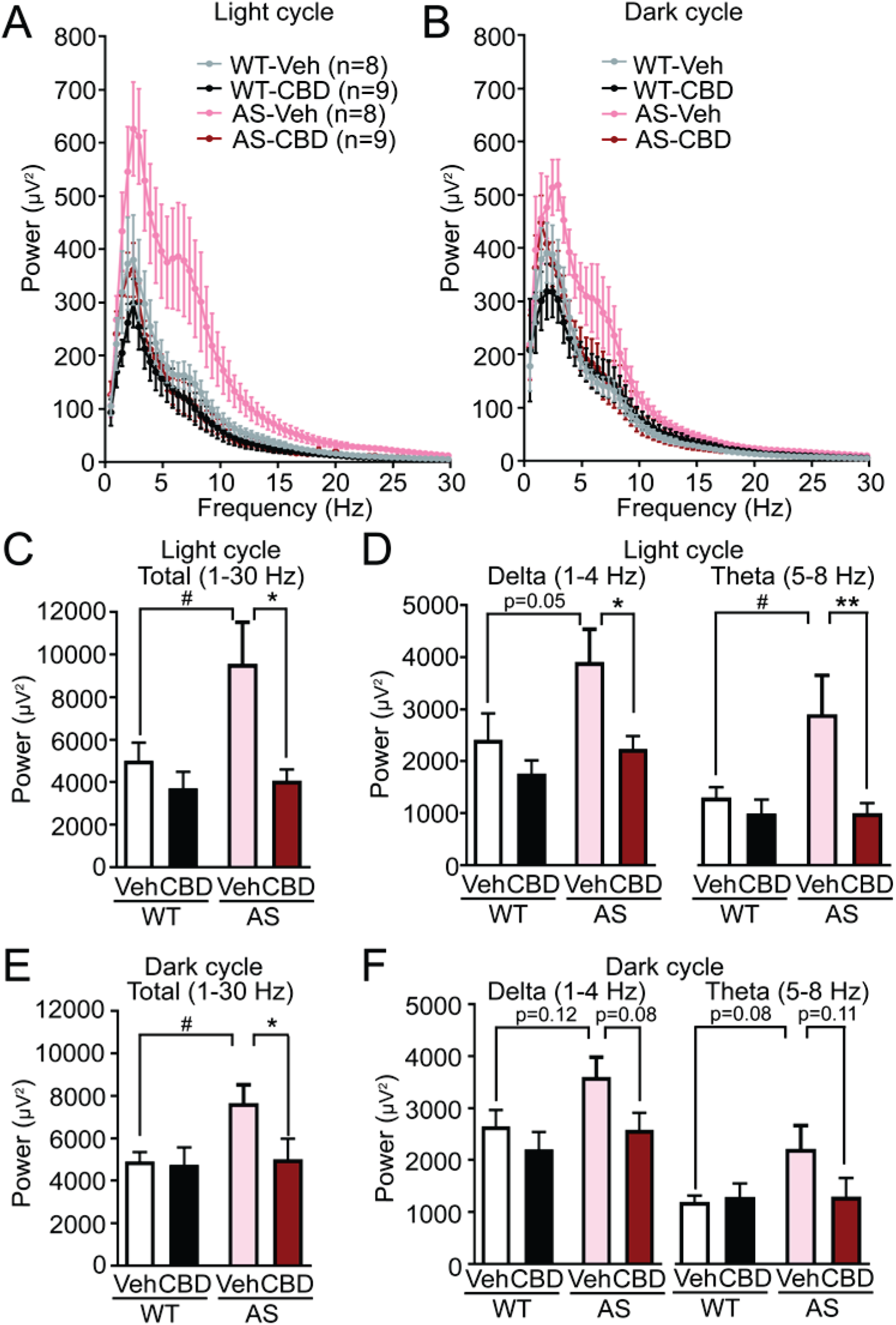
Two-weeks of CBD administrations normalize LFP in AS model mice. (A and B) Power analyses of cortical LFP during (A) light or (B) dark cycle of WT and AS model mice treated with Veh or CBD (100 mg/kg, once daily, i.p.). (C–F) LFP power analyses of (C and E) total (1–30 Hz), as well as (D and F) delta δ (1–4 Hz) and theta θ (5–8 Hz) frequency bands during (C and D) light or (E and F) dark cycle of WT and AS model mice treated with Veh or CBD (100 mg/kg, once daily, i.p.). n=8–9 mice/group. # p<0.05 compared to WT-Veh; * p<0.05, **p<0.01 compared to AS-Veh, two-way ANOVA with Tukey’s *post hoc* test.

Our data are the first to show that CBD can reduce both acoustically- and hypertherimia-induced seizure severity in AS model mice. As such, our findings suggest that CBD might attenuate seizures in AS individuals, which expands the therapeutic spectrum of the anti-epileptic effects of CBD (17, 19, 23, 27-32). The fact that neither acute nor chronic post-kindling CBD treatment affected flurothyl- or hyperthermia-induced seizure threshold in AS model mice suggests that further evaluation of CBD using additional models of epilepsy is required to reveal the full antiepileptic potential of CBD in AS. The context-dependent antiepileptic effects of CBD suggest that different seizure models can engage distinct mechanisms for seizure initiation and propagation. And, relevant to human use, such a finding indicates that CBD might be beneficial for some types of seizures and not others, depending on the circuits engaged and/or at what stage of epilepsy intervention begins.

CBD shows behavioral benefits in animal models of motor, social, and cognitive impairments (33-37). However, the behavioral benefits of CBD in animal models often exhibit an inverted U-shaped dose-response curve, with higher doses (> 20 mg/kg) being ineffective and sedative, whereas lower doses can prove effective (38, 39). This conundrum for designing CBD dosing that both control seizures and improve behavior was also found in studies of Dravet syndrome model mice (40). The same may be true for AS, as the effective concentration (e.g. 100 mg/kg) at which we observed anticonvulsant effects also produced sedative effects. Notably, the sedative effects in mice at higher doses of CBD preclude meaningful interpretation of many movement dependent behavioral paradigms (e.g. 3 chamber sociability test, fear-conditioning, and Morris water maze), and may also explain the reduced marble burying behavior, which is associated with locomotor activity (41). However, while AS model mice recapitulate many AS-like behaviors, activity is not one of those; AS model mice are hypoactive, while AS individuals are often hyperactive (42). Thus, while anticonvulsant doses of CBD (i.e. 100 mg/kg) are mildly sedative in AS model mice, it is conceivable that analogous CBD dosing could alleviate hypermotor behavior often observed in AS individuals.

EEG power is broadly increased in children with AS relative to age-matched neurotypical controls, and this is recapitulated under certain conditions in AS model mice relative to WT littermates (7, 24). Among the elevated EEG spectrum, enhanced delta rhythm is a particularly reliable biomarker for AS (13). Here we found that two weeks of CBD administrations reduced wide spectral cortical electrophysiological activities, and normalized the delta (1–4 Hz) and theta (5–8 Hz) activity in AS model mice. This suggests that CBD can suppress pathological EEG signatures in AS model mice, and perhaps in AS individuals.

This study lends critical support for using CBD to treat seizures, along with behavioral and EEG abnormalities in AS, and expands the potential beneficial spectrum of CBD.

## METHODS

Detailed experimental methods are included with the supplemental materials. Male and female mice were used for experiments in equal genotypic ratios. Synthetic CBD (99.2±0.18% purity) was provided by RTI International (Log #3857-52-1). Data are presented as mean ± SEM. Unless otherwise noted, unpaired t-test (two-tailed) were used for single comparisons; 2-way ANOVAs were used for multiple comparisons. P value < 0.05 was considered significant.

### Study approval

All animal procedures followed NIH guidelines and were approved by the IACUC at the University of North Carolina.

## Author contributions

BG, MZ, and MRG performed experiments. BG, MZ, and MRG analyzed the data. MR managed the mouse colony and genotyping. BG, VDN, SSM, PRC, and BDP designed and coordinated the investigations. BG, PRC, and BDP wrote the manuscript.

## Acknowledgments

This work was supported by the Angelman Syndrome Foundation (to PRC and BDP), NIH grant R01HD093771 (to BDP), and NICHD U54 Intellectual and Developmental Disabilities Research Center HD079124 (to SSM).

## Notes

The authors have declared that no conflicts of interest exist.

## REFERENCES

1. Buiting K, Williams C, and Horsthemke B. Angelman syndrome - insights into a rare neurogenetic disorder. Nature reviews Neurology. 2016;12(10):584–93.

2. Williams CA, Driscoll DJ, and Dagli AI. Clinical and genetic aspects of Angelman syndrome. Genetics in medicine: official journal of the American College of Medical Genetics. 2010;12(7):385–95.

3. Bakke KA, Howlin P, Retterstol L, Kanavin OJ, Heiberg A, and Naerland T. Effect of epilepsy on autism symptoms in Angelman syndrome. Molecular autism. 2018;9:2.

4. Fiumara A, Pittala A, Cocuzza M, and Sorge G. Epilepsy in patients with Angelman syndrome. Italian journal of pediatrics. 2010;36:31.

5. Laan LA, et al. Evolution of epilepsy and EEG findings in Angelman syndrome. Epilepsia. 1997;38(2):195–9.

6. Thibert RL, et al. Epilepsy in Angelman syndrome: a questionnaire-based assessment of the natural history and current treatment options. Epilepsia. 2009;50(11):2369–76.

7. Born HA, et al. Strain-dependence of the Angelman Syndrome phenotypes in Ube3a maternal deficiency mice. Scientific reports. 2017;7(1):8451.

8. Gu B, et al. Ube3a reinstatement mitigates epileptogenesis in Angelman syndrome model mice. The Journal of clinical investigation. 2018.

9. Jiang YH, et al. Mutation of the Angelman ubiquitin ligase in mice causes increased cytoplasmic p53 and deficits of contextual learning and long-term potentiation. Neuron. 1998;21(4):799–811.

10. Judson MC, et al. GABAergic Neuron-Specific Loss of Ube3a Causes Angelman Syndrome-Like EEG Abnormalities and Enhances Seizure Susceptibility. Neuron. 2016;90(1):56–69.

11. Mandel-Brehm C, Salogiannis J, Dhamne SC, Rotenberg A, and Greenberg ME. Seizure-like activity in a juvenile Angelman syndrome mouse model is attenuated by reducing Arc expression. Proc Natl Acad Sci U S A. 2015;112(16):5129–34.

12. Miura K, et al. Neurobehavioral and electroencephalographic abnormalities in Ube3a maternal-deficient mice. Neurobiology of disease. 2002;9(2):149–59.

13. Sidorov MS, et al. Delta rhythmicity is a reliable EEG biomarker in Angelman syndrome: a parallel mouse and human analysis. Journal of Neurodevelopmental Disorders. 2017;9(1):17.

14. Silva-Santos S, et al. Ube3a reinstatement identifies distinct developmental windows in a murine Angelman syndrome model. The Journal of clinical investigation. 2015.

15. Sonzogni M, et al. A behavioral test battery for mouse models of Angelman syndrome: a powerful tool for testing drugs and novel Ube3a mutants. Molecular autism. 2018;9:47.

16. Izzo AA, Borrelli F, Capasso R, Di Marzo V, and Mechoulam R. Non-psychotropic plant cannabinoids: new therapeutic opportunities from an ancient herb. Trends Pharmacol Sci. 2009;30(10):515–27.

17. Devinsky O, et al. Open-label use of highly purified CBD (Epidiolex(R)) in patients with CDKL5 deficiency disorder and Aicardi, Dup15q, and Doose syndromes. Epilepsy Behav. 2018;86:131–7.

18. Hess EJ, et al. Cannabidiol as a new treatment for drug-resistant epilepsy in tuberous sclerosis complex. Epilepsia. 2016;57(10):1617–24.

19. Devinsky O, et al. Cannabidiol in patients with treatment-resistant epilepsy: an open-label interventional trial. Lancet Neurol. 2016;15(3):270–8.

20. Gofshteyn JS, et al. Cannabidiol as a Potential Treatment for Febrile Infection-Related Epilepsy Syndrome (FIRES) in the Acute and Chronic Phases. Journal of child neurology. 2017;32(1):35–40.

21. Ibeas Bih C, Chen T, Nunn AV, Bazelot M, Dallas M, and Whalley BJ. Molecular Targets of Cannabidiol in Neurological Disorders. Neurotherapeutics. 2015;12(4):699–730.

22. Huang HS, et al. Behavioral deficits in an Angelman syndrome model: effects of genetic background and age. Behavioural brain research. 2013;243:79–90.

23. Jones NA, et al. Cannabidiol exerts anti-convulsant effects in animal models of temporal lobe and partial seizures. Seizure. 2012;21(5):344–52.

24. Frohlich J, et al. Electrophysiological Phenotype in Angelman Syndrome Differs Between Genotypes. Biological psychiatry. 2019;85(9):752–9.

25. Valente KD, et al. Angelman syndrome: difficulties in EEG pattern recognition and possible misinterpretations. Epilepsia. 2003;44(8):1051–63.

26. Vendrame M, et al. Analysis of EEG patterns and genotypes in patients with Angelman syndrome. Epilepsy Behav. 2012;23(3):261–5.

27. Devinsky O, et al. Trial of Cannabidiol for Drug-Resistant Seizures in the Dravet Syndrome. N Engl J Med. 2017;376(21):2011–20.

28. Devinsky O, et al. Long-term cannabidiol treatment in patients with Dravet syndrome: An open-label extension trial. Epilepsia. 0(0).

29. Filloux FM. Cannabinoids for pediatric epilepsy? Up in smoke or real science? Transl Pediatr. 2015;4(4):271–82.

30. Friedman D, and Devinsky O. Cannabinoids in the Treatment of Epilepsy. N Engl J Med. 2015;373(11):1048–58.

31. Rosenberg EC, Tsien RW, Whalley BJ, and Devinsky O. Cannabinoids and Epilepsy. Neurotherapeutics. 2015;12(4):747–68.

32. Verrotti A, Castagnino M, Maccarrone M, and Fezza F. Plant-Derived and Endogenous Cannabinoids in Epilepsy. Clin Drug Investig. 2016;36(5):331–40.

33. Almeida V, et al. Cannabidiol exhibits anxiolytic but not antipsychotic property evaluated in the social interaction test. Progress in neuro-psychopharmacology & biological psychiatry. 2013;41:30–5.

34. Campos AC, Fogaca MV, Sonego AB, and Guimaraes FS. Cannabidiol, neuroprotection and neuropsychiatric disorders. Pharmacological research. 2016;112:119–27.

35. Gururajan A, Taylor DA, and Malone DT. Cannabidiol and clozapine reverse MK-801-induced deficits in social interaction and hyperactivity in Sprague-Dawley rats. J Psychopharmacol. 2012;26(10):1317–32.

36. Malone DT, Jongejan D, and Taylor DA. Cannabidiol reverses the reduction in social interaction produced by low dose Delta(9)-tetrahydrocannabinol in rats. Pharmacology, biochemistry, and behavior. 2009;93(2):91–6.

37. Peres FF, et al. Cannabidiol Prevents Motor and Cognitive Impairments Induced by Reserpine in Rats. Front Pharmacol. 2016;7:343.

38. Campos AC, Moreira FA, Gomes FV, Del Bel EA, and Guimaraes FS. Multiple mechanisms involved in the large-spectrum therapeutic potential of cannabidiol in psychiatric disorders. Philos Trans R Soc Lond B Biol Sci. 2012;367(1607):3364–78.

39. Guimaraes FS, Chiaretti TM, Graeff FG, and Zuardi AW. Antianxiety effect of cannabidiol in the elevated plus-maze. Psychopharmacology. 1990;100(4):558–9.

40. Kaplan JS, Stella N, Catterall WA, and Westenbroek RE. Cannabidiol attenuates seizures and social deficits in a mouse model of Dravet syndrome. Proc Natl Acad Sci U S A. 2017;114(42):11229–34.

41. Kim H, Lim CS, and Kaang BK. Neuronal mechanisms and circuits underlying repetitive behaviors in mouse models of autism spectrum disorder. Behavioral and brain functions: BBF. 2016;12(1):3.

42. Pelc K, Cheron G, and Dan B. Behavior and neuropsychiatric manifestations in Angelman syndrome. Neuropsychiatric disease and treatment. 2008;4(3):577–84.

