## Supplemental Materials for "Cannabidiol attenuates seizures and EEG abnormalities in Angelman syndrome model mice"

### Supplemental Methods

#### Mice

All animal procedures were approved by the Institutional Animal Care and Use Committee of the University of North Carolina at Chapel Hill and were performed in accordance with the guidelines of the U.S. National Institutes of Health. Mice were group-housed on a 12:12 light/dark cycle with *ad libitum* access to food and water. Male and female mice were used for experiments in equal genotypic ratios. Mice were randomly assigned to experimental groups. Our AS mouse colony has been maintained on two different congenic background strains: 129S2/SvPasCrl (129) and C57BL/6J (B6). Dr. Ype Elgersma (Erasmus Medical Center) provided the 129 mice. B6 mice were purchased from Jackson Labs (JAX #: 016590). AS model mice ($Ube3a^{m–/p+}$) and their wild-type (WT) littermates were generated by breeding $Ube3a^{m+/p–}$ females with WT males.

#### Drug treatment

Synthetic CBD (99.2±0.18% purity) was provided by RTI International (Log #3857-52-1). CBD was freshly prepared by dissolving in the solvent of ethanol, cremophor, and 0.9% saline at the ratio of 2:1:17. CBD or Vehicle (equal amount of solvent) was intraperitoneally (i.p.) injected at the indicated doses.

**Audiogenic-induced seizure**

We tested P70–P100 mice that were backcrossed over 10 generations onto the 129 background to maximize the penetrance of audiogenic seizure phenotypes. Following 1 minute of habituation to the behavioral chamber, we exposed mice to 1 minute of alarm sirens (~125 dB; 49-1010, RadioShack, Fort Worth, TX). We video-recorded each session and scored seizure severity blind to genotype and treatment. The response of each mouse was scored on the basis of the most severe phenomenon seen (i.e., no seizure response = 0; wild running and jumping = 1; tonic-clonic seizure = 2), and a mean seizure severity score was calculated. Veh or CBD was injected (10, 20, 50 or 100 mg/kg, i.p.) 1 hr prior to the start of testing.

**Flurothyl kindling**

For flurothyl induced seizures, each B6 mouse was allowed to habituate for 1 min in a 2 liter glass chamber before the top of the chamber was closed. Then, 10% flurothyl (bis-2,2,2-trifluoroethyl ether; Sigma-Aldrich) in 95% ethanol was infused at a rate of 200 µL/min onto a disk of filter paper (Whatman, Grade 1) suspended at the top of the chamber. We then recorded duration to myoclonic seizure (sudden involuntary jerk/shock-like movements involving the face, trunk, and/or limbs) and generalized seizure (also known as clonic-forebrain seizures that are characterized by clonus of the face and limbs, loss of postural control, rearing, and falling). Upon emergence of a generalized seizure, the lid of the chamber was immediately removed, allowing for rapid dissipation of the flurothyl vapors and exposure of the mouse to fresh air. Mice were then returned to their home cage following recovery from behavioral seizures. The flurothyl chamber was recharged with fresh filter paper, cleaned using water, and thoroughly dried between subjects.

For kindling, flurothyl exposures were repeated once daily over eight consecutive days (induction phase). Mice were then given a 28-day rest period (incubation phase), during which they remained in their home cages with no flurothyl exposure, and then re-exposed to flurothyl once more on day 36 (retest). For chronic treatment, Veh or CBD administrations were initiated immediately after mice recovered from flurothyl-seizure on day 8 and continued for 2 weeks (100 mg/kg, i.p., once per day), and then flurothyl retest was performed after 2 weeks of drug washout. For acute treatment, Veh or CBD (100 mg/kg, i.p.) was injected 1 hr prior to the flurothyl retest. Mouse behavior during each flurothyl exposure was video-recorded and reviewed by investigators, blind to both genotype and treatment, who determined latency to the onset of myoclonic and generalized seizures.

**Hyperthermia-induced seizures in kindled AS model mice**

A week after flurothyl retest, core body temperature was monitored by rectal probe and maintained at a command temperature ± 0.1°C using a rodent temperature controller (TCAT-2DF, Physitemp Inc.) and heat lamp positioned above the mouse. Mice were first held at 36.5°C for 10 min before the body temperature was increased incrementally by 0.3°C per min until the emergence of behavioral seizures scored 3 and above based on a modified Racine Scale (described below) or until a core body temperature of 42.5°C was reached. For each mouse, the duration and severity of seizures were determined from video recordings. Seizure severity was assessed based on a modified Racine Scale scoring system: 1 = mouth and facial movements; 2 = head nodding; 3 = forelimb clonus, usually one limb; 4 = forelimb clonus with rearing; 5 = generalized whole body clonus, rearing and falling; and 6 = generalized seizures with tonic muscle extension of limbs and trunk. Because seizures with a severity below 3 are difficult to assess in mice (1), we limited our seizure severity assessment to seizures with a Racine score of 3-6. Veh or CBD (100 mg/kg, i.p.) was injected 1 hr prior to the start of testing.

**Open field test**

Locomotor activity of adult B6 mice in a novel environment was assessed by 1 hr trial in a photocell-equipped automated chamber (41 cm × 41 cm × 30 cm; Versamax system, Accuscan Instruments). Measures were taken of total distance traveled and number of rearing movements. Activity chambers were contained inside sound-attenuating boxes equipped with ceiling-mounted lights and fans. Veh or CBD was injected (10, 50 or 100 mg/kg, i.p.) 1 hr prior to the start of testing.

**Rotarod performance**

Adult B6 mice were assessed for balance and motor coordination on an accelerating rotarod (Ugo-Basile, Stoelting Co., Wood Dale, Il). Revolutions per minute (rpm) were set at an initial value of 3, with a progressive increase to a maximum of 30 rpm across 5 min, the maximum trial length. Test sessions consisted of 3 trials of acquisition session and 2 trials of retest session 48 hr later, with 45 sec between each trial. Latency to fall was measured by the rotarod timer. Veh or CBD (10, 50 or 100 mg/kg, i.p.) was injected 1 hr prior to the start of acquisition session.

**Marble-burying assay**

Adult B6 mice were tested in a Plexiglas cage located in a sound-attenuating chamber with ceiling light and fan. The cage contained corncob bedding 5 cm deep, with 20 black glass marbles (14 mm diameter) arranged in an equidistant 5 × 4 grid on top of the bedding. Animals were given access to the marbles for 30 min. An overhead photograph was taken of the cage at the start and end of the session, and then the image was converted to grayscale and analyzed using ImageJ. Using the uncovered marble as a baseline, the image of the interior of the cage was thresholded and the particle analyze function was used to count the number of marbles in which greater than 33% of the area was covered with bedding. Veh or CBD (10, 50 or 100 mg/kg, i.p.) was injected 1 hr prior to the start of testing.

**Local field potential (LFP) recordings in freely moving mice**

For surgeries, adult (P60–P90) B6 mice were anesthetized via intraperitoneal injections of ketamine (40 mg/kg) and xylazine (10 mg/kg), and 0.25% bupivacaine was applied topically for local analgesia. Stainless steel bipolar recording electrodes (Plastics One Inc.) were implanted in right motor cortex (coordinates from bregma: AP=-1.0 mm; L=1.0 mm; and D=-0.5 mm below dura), while ground electrodes were fastened to a stainless steel screw positioned on the skull above the cerebellum. Electrode positions were secured using dental cement. Mice were allowed to recover for 7 days before recording. A tethered system with a commutator (Plastics One Inc.) was used for recordings, allowing mice to roam freely within the home cage and enabling time-locked video-LFP recording of mouse brain function in conjunction with behavior. Mice were recorded for 24 hr prior to (baseline activity) and after 2-week Veh or CBD treatment (100 mg/kg, i.p., once per day). LFP recordings were amplified (1,000x) using single-channel amplifiers (Grass Technologies), sampled at a rate of 1000 Hz, and filtered with 0.3 Hz high-pass and 100 Hz low-pass filters. All electrical data were digitized with CED Micro1401 (Cambridge Electronic Design Ltd.). We used Spike2 software (Cambridge Electronic Design Ltd.) for spectral analysis of LFP samples during the full hour of recording between 12 pm to 1 pm as a representation of activity during the light cycle and 12 am to 1 am as a representation of activity during the dark cycle. We analyzed the total spectral power (1–30 Hz), as well as delta δ (1–4 Hz) and theta θ (5–8 Hz) frequency bands. To make clear the analogy to human studies, LFP recordings in mice are sometimes referred to in the text as EEG recordings.

**Statistics**

All experiments and analyses were performed blind to genotype and treatment, and all statistical analyses were performed using GraphPad Prism 7.04 software (GraphPad Software Inc.). Unless otherwise noted, comparisons between two groups were analyzed using unpaired t-tests (two-tailed), while multi-group comparisons were analyzed using 2-way ANOVA with Bonferroni's or Tukey's *post hoc* test. $P < 0.05$ was considered significant.

### Supplemental References

1. Kaplan JS, Stella N, Catterall WA, and Westenbroek RE. Cannabidiol attenuates seizures and social deficits in a mouse model of Dravet syndrome. *Proc Natl Acad Sci U S A.* 2017;114(42):11229-34.

### Supplemental Figures

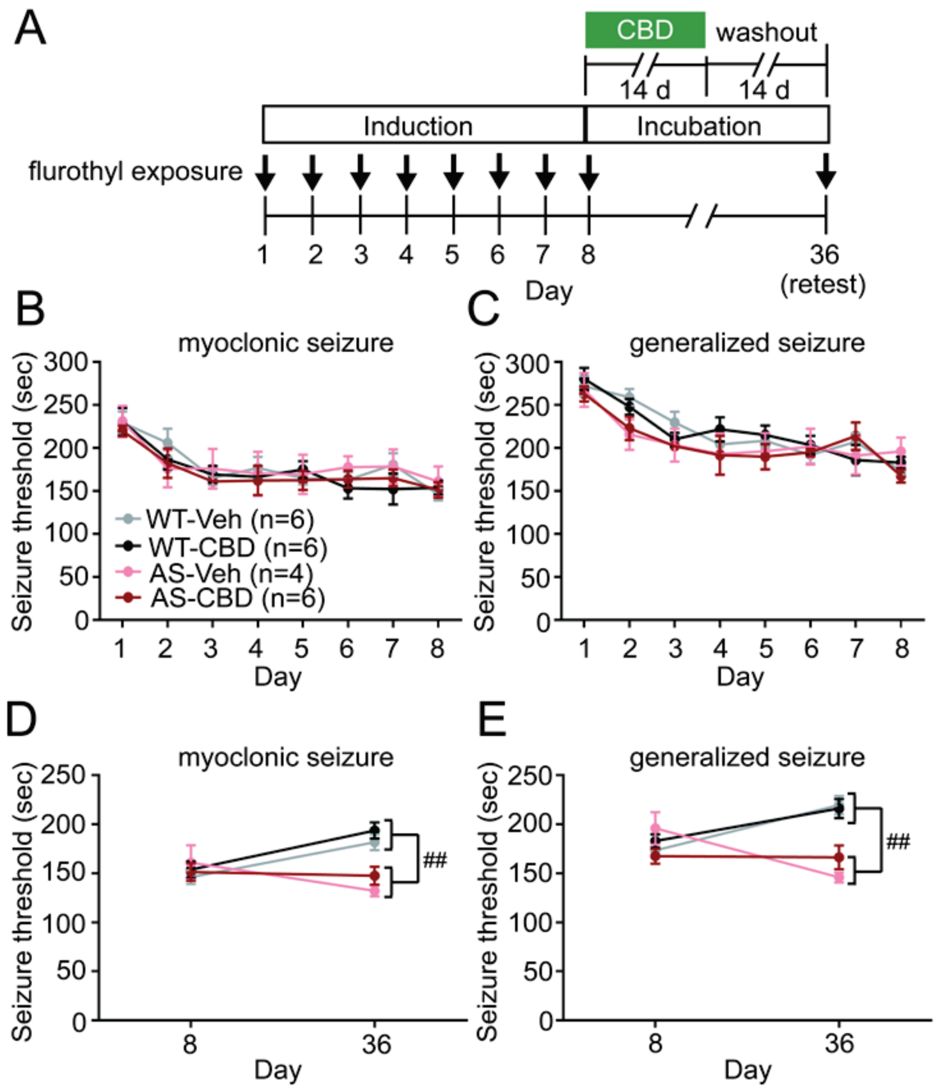


**Supplemental Figure 1. A two-week treatment of CBD after flurothyl induction fails to prevent elevated retest seizure susceptibility in AS model mice.** (A) Schematic of flurothyl kindling and CBD treatment. (B) Myoclonic and (C) generalized seizure threshold during 8-day flurothyl induction of WT and AS model mice prior to initiation of treatment. (D) Retest myoclonic and (E) generalized seizure threshold of WT or AS model mice treated with Veh or CBD (100 mg/kg, i.p.) during the first two weeks of incubation period. n=4-6 mice/group. ## $p<0.01$ AS compared to WT, two-way ANOVA.

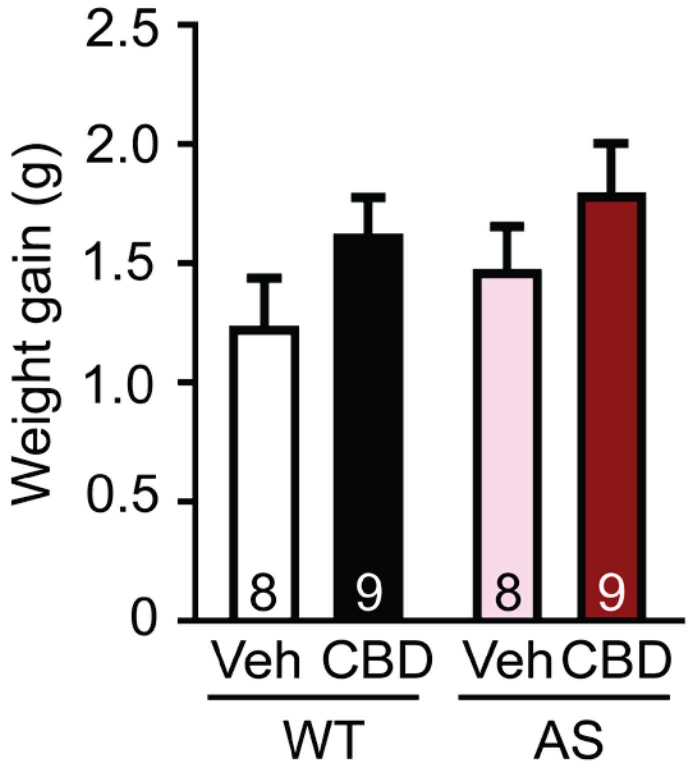


**Supplemental Figure 2. A two-week treatment of CBD did not affect weight gain in WT and AS model mice.** Body weight was measured prior to and after 2-week treatment of Veh or CBD in the cohort of WT and AS model mice used for LFP recording and gain of body weight was calculated. n=8-9 mice/group. No significant difference was found using two-way ANOVA analysis.
